# Environmental DNA identifies individual free-ranging tiger sharks from a multi-use nuclear genotyping panel

**DOI:** 10.64898/2026.11.22.768475

**Authors:** Jessica R. Pearce, Abinaya Meenakshisundaram, Oliver Jewell, Ben D’Antonio, Luciana C. Ferreira, Mark G. Meekan, Shannon Corrigan, Luke Thomas

## Abstract

Conventional monitoring of marine wildlife, including tagging and tissue sampling for population genetic analyses, can be invasive, resource-intensive, and time-consuming. For marine organisms, targeted seawater environmental DNA (eDNA) sampling is emerging as a non-invasive alternative to traditional tissue biopsies, but recovering nuclear DNA and genotyping individuals from environmental samples remains challenging. Microhaplotypes, short DNA fragments containing multiple variant sites, can provide high discriminatory power for individual identification while enabling population genetic analyses that account for linkage disequilibrium in low-input or degraded samples. Here, we present the first microhaplotype panel for the tiger shark (*Galeocerdo cuvier*) and evaluate its application across conventional tissue and non-invasive eDNA sampling from sharks in Western Australia. Our 38-marker panel, comprising 151 microhaplotypes, discriminated individual sharks with high accuracy, as paired eDNA and tissue samples showed consistent clustering. Estimates of genetic variation using eDNA derived genotypes were representative of whole genome data and revealed a largely panmictic population along the coastline of Western Australia. Importantly, microhaplotypes recovered from non-targeted water samples correctly detected a shark present in the area confirmed by acoustic detections. Together, these results demonstrate the potential of PCR-based genotyping approached for cost-effective population genetic analyses and individual identification from tissue and degraded environmental samples, providing a significant step towards scalable, non-invasive monitoring of wild shark populations.

## 1. INTRODUCTION

In the current era of rapid biodiversity loss due to anthropogenic change, scalable and routine wildlife population monitoring is important to inform appropriate conservation and management measures (Ceballos et al., 2015; Cowie et al., 2022). Conventional approaches to studying movement include visual surveys, acoustic and satellite telemetry, and assessments of genetic connectivity, providing complementary information on individual behaviour and population genetic characteristics (Ferreira et al., 2018; Hohenlohe et al., 2021; Matley et al., 2022). However, these approaches require visual census and often animal capture, and can be prohibitively invasive, costly or resource-intensive, limiting their spatial and temporal scalability. Non-invasive, cost-effective alternatives are urgently needed to enable broader-scale monitoring (Carroll et al., 2018).

Environmental DNA (eDNA), comprising trace DNA shed by organisms into their environment, offers a promising alternative for wildlife monitoring (Altermatt et al., 2025; Blackman et al., 2024). eDNA metabarcoding is now well established for broadscale biodiversity assessment (Miya, 2022; Pearce, Ayad, et al., 2026; Stat et al., 2017; West et al., 2021; West et al., 2020), and recent advances have demonstrated its potential for making population genetic inferences and individual identification (Altermatt et al., 2025; Jensen et al., 2021; Sigsgaard et al., 2020). Mitochondrial DNA (mtDNA) markers can recover intraspecific variation from environmental samples (Dugal et al., 2022) but are maternally inherited and contain insufficient variation for comprehensive individual genotyping. Nuclear DNA (nuDNA) markers, particularly single nucleotide polymorphisms (SNPs), therefore offer greater discriminatory power (Andres et al., 2023; Jensen et al., 2021). However, nuDNA is less abundant and presumably less persistent in the environment than mtDNA, making its recovery and reliable genotyping from environmental samples challenging (Andres et al., 2021; Jensen et al., 2021). Individual identification of free-ranging animals from environmental samples (e.g. scat) has been successfully demonstrated in terrestrial mammals, including bears, big cats and wolves (De Barba et al., 2024; Solari et al., 2024). The individual identification of fish from aquaria water samples has likewise been demonstrated (Meenakshisundaram, Jarman, et al., 2025), although extension to in situ detections in aquatic environments remains in development (Jensen et al., 2021; Meenakshisundaram, Meekan, et al., 2025; Parsons et al., 2018). Microhaplotypes offer a promising marker choice as they contain multiple SNPs, providing higher discriminatory power within relatively short amplicons (<300 bp) (Anderson et al., 2025; Baetscher et al., 2018; Meenakshisundaram, Jarman, et al., 2025; Turchi et al., 2025).

The tiger shark, *Galeocerdo cuvier* (Péron & Lesueur, 1822), is a charismatic, cosmopolitan species distributed throughout tropical and warm-temperate waters (Last & Stevens, 2009). Through their predatory role, tiger sharks can influence the structure and functioning of marine ecosystems, although they face substantial, compounding threats including fishing and habitat degradation (Braccini et al., 2021; Ferreira et al., 2017; Heithaus et al., 2012). Their extensive movements (Ferreira et al., 2015; Ostrovski et al., 2026) and use of diverse marine habitats (D’Antonio et al., 2025; Lipscombe et al., 2020; Werry et al., 2014) present challenges for monitoring populations across broad spatial and temporal scales. Developing scalable monitoring approaches is therefore important for improving our understanding of the distribution, movements and population dynamics of highly mobile marine species. Here, we present a microhaplotype panel designed for tiger sharks and evaluate its application for individual detections based on conventional population genetic and environmental sampling. We demonstrate its power for identifying individuals by matching genetic profiles between paired tissue and eDNA samples from the same shark and highlight its potential scalability and various applications. Our results support the development of nuclear DNA-based PCR genotyping approaches for wildlife monitoring and management using sampling of environmental DNA.

## 2. METHODS

### 2.1 Sample collection

Field trips were conducted from 2023 to 2025 to tag and sample tiger sharks from the Exmouth Gulf (*n* = 17) and the Mid-West region of Western Australia (*n* = 4) (Figure 1). Sharks were caught using baited drumlines and brought alongside the boat. While the shark was restrained next to boat, we collected targeted eDNA samples as four replicate 2L seawater samples from the surface at the stern of the vessel. In addition to these eDNA samples, small tissue fin clips were taken with a scalpel from the pectoral fin of each shark and flash frozen in liquid nitrogen. Samples were stored at −80℃ upon return to laboratory facilities. Paired tissue (*n* = 21) and target eDNA samples (*n* = 84) were taken from a total of 21 individual tiger sharks (19 females; 2 males) ranging from 205-375cm total length spanning 11 sites across the three years (Figure 1). We also surgically inserted internal acoustic tags into the abdominal cavity of each shark to track their movements across acoustic receiver arrays in these areas (as described in Bruce et al., 2019) as a complementary detection method that could be used to validate eDNA-based detections. Water sampling was always conducted before surgery to avoid blood confounding the eDNA results.

**Figure 1.**
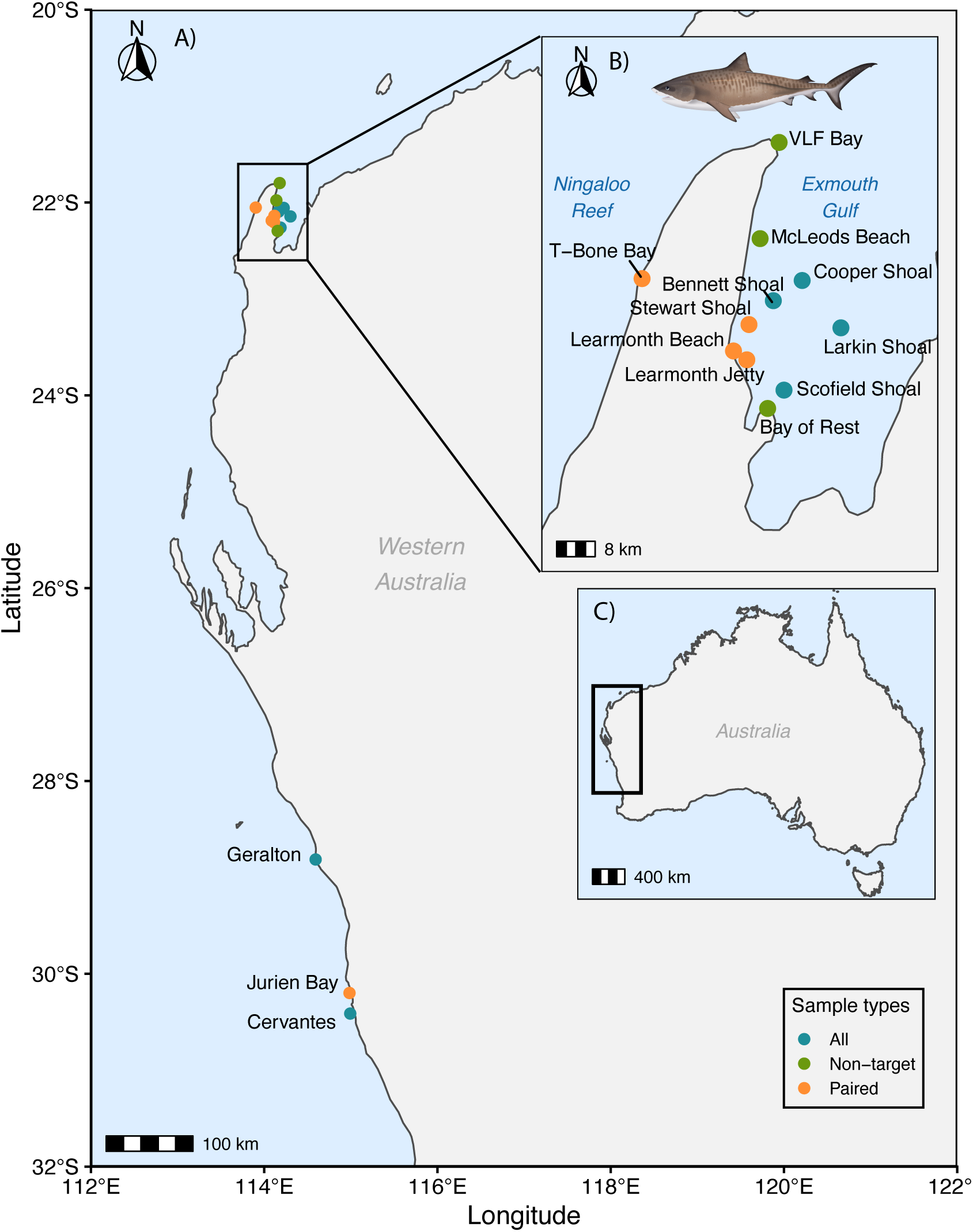
A) Sampling locations along the Western Australian coastline. Paired sampling = tissue and targeted eDNA from same tiger shark individual. Non-target = only eDNA samples without tiger sharks present B) Sample locations from the Gascoyne region. C) Black box denotes the position of the sampling coastline in Australia. Tiger shark image ©MarineWise.com.au.

We also collected ‘non-targeted’ seawater samples from a subset of sites. This sampling was conducted opportunistically when logistics allowed at a minimum of 100 m distance from any targeted eDNA sampling or tagging that had occurred that day, using freshly sterilised collection bottles and gloves, following the same 4 replicates of 2L design (*n sites* = 11; *n samples* = 43). For both target and non-target eDNA samples, samples were stored on ice until filtration and replicates were filtered separately through a 0.45 µm mixed-cellulose ester filter membrane using a Masterflex peristaltic pump within 24 hours post-collection. Filter papers were immediately placed into 2 mL DNA LoBind tubes (Eppendorf) containing 540 µL Buffer ATL (Qiagen) and tubes were stored at −20°C until return to laboratory facilities. Filtration equipment was sterilised in 10% commercial-grade bleach to prevent cross-contamination and rinsed with deionised water between replicates and sites. Two litres of the rinse water were also filtered after each site as a negative control to account for potential contamination at this stage and assess sterilisation effectiveness.

### 2.2 Tiger shark microhaplotype panel

#### 2.2.1 Samples and data generation for panel design

The microhaplotype panel created for this study was designed from population-level whole genome sequencing (WGS) data. In brief, DNA from 130 tiger shark samples collected from south-west to northern Australia and South Africa were extracted using DNeasy® Blood and Tissue Kit (Qiagen) standard protocol. The Illumina DNA Prep kit was used for library preparation and then samples were sequenced at roughly 10X depth on an Illumina NovaSeq 6000 S4 flowcell. This dataset also includes four samples that were previously processed and published with the same methodology but sequenced at higher (∼35X) coverage (Pearce, Doran, et al., 2026). Fastp quality trimmed reads (Chen et al., 2018) were mapped to Haplotype 1 of the tiger shark reference genome (Genbank accession: GCA_062075035.1; (Pearce, Doran, et al., 2026) with bwa-mem2 v2.2.1 (Vasimuddin et al., 2019). Samtools v1.16.1 (Danecek et al., 2021) was then used to filter, sort and index the resulting bam files (--min-MQ 10). Duplicate reads were removed with the MarkDuplicates function (--REMOVE_DUPLICATES true) from the Picard package within GATK4 v4.2.5.0 (McKenna et al., 2010). Variants were called and filtered using a combination of BCFtools (Danecek et al., 2021) and VCFtools (Danecek et al., 2011) to retain only biallelic SNPs with a minimum of 10X coverage, a minor allele frequency of 0.05 and heterozygosity between 0.18 and 0.5.

#### 2.2.2 Panel development

To begin identifying candidate microhaplotypes, we extracted all SNPs that passed the previous filters using BCFtools *view*. BEDtools *cluster* function (Quinlan, 2014) was then employed to find 250 base pair (bp) regions that contained multiple SNPs (Podini et al., 2026). These were filtered to only retain those with at least 3 SNPs per region. The corresponding fasta sequences were extracted for the resulting candidate microhaplotype regions with BEDtools *getfasta* function. These candidate microhaplotype sequences were locally aligned using BLAST v2.17.0 (Madden, 2013) against the reference assembly (-evalue 1e-3-perc_identity 100 - qcov_hsp_perc 100) to check for sequence specificity, and only sequences that mapped to one location on the genome were retained. This process was repeated to check for species specificity by blasting against the NCBI nt database instead and removing any sequences that mapped to other species (evalue 1e-3-perc_identity 90 - qcov_hsp_perc 100).

Primers were designed for these final microhaplotype sequences using NGS-PrimerPlex (Kechin et al., 2020) with the following options: primer length: 18-80bp, amplicon length: 200-260bp (optimal = 250bp), primer melt: 50-60℃ (optimal = 55℃), primer GC content: 40-60% (optimal = 50%), and with the blast option on. Following primer generation, we repeated the same previous checks for sequence and species-specificity process for the primer sequences though BLAST. Any primer sequences (forward or reverse) were removed if they had a BLAST hit using the previously stated parameters against another species or multiple locations within the reference assembly, then only intact primer pairs were retained. We rechecked the internal amplicons produced by NGS-PrimerPlex for sequence and species specificity to ensure confidence in the final panel. A final panel of 80 microhaplotype primer pairs were selected and synthesised through Integrated DNA Technologies (IDT) with added Illumina overhang adapter sequences (Supplementary materials 2). Once received, separate PCR reactions were conducted using each primer pair to test amplification efficacy on tiger shark DNA derived from tissue samples. Primer pairs that failed to amplify targets effectively, or those that amplified off-target DNA, were not taken forward for the final microhaplotype multiplex panel (Supplementary materials 2).

#### 2.2.3 Sample laboratory processing

Genomic DNA (gDNA) from all tissue samples was extracted with the DNeasy® Blood and Tissue Kit (Qiagen) standard protocol with a 3-hour digestion. Extracted gDNA concentration was tested with the Qubit dsDNA BR (Broad Range) kit on a Qubit 4 Fluorometer (Thermo Fisher Scientific Inc., Waltham, MA, USA). Environmental DNA extractions were performed using the DNeasy® Blood and Tissue Kit (Qiagen), with modifications to the standard protocol to improve DNA yield (Missen, Ayad, et al., 2026). Negative controls (extraction blanks) were processed with each batch of extractions to detect any possible laboratory contamination during extraction.

Multiplex primers that passed the PCR efficacy test (successfully amplified on-target and with no off-target amplification) were combined into a working stock containing 0.4 µM per primer pair (0.2 µM forward and reverse). The first-stage multiplex PCR was conducted with a final primer concentration of 0.08 µM using 2 ng DNA input for gDNA samples and 5 µl of extracted DNA for eDNA samples which varied a great deal in concentration (Table S1) with a two-step touch-down PCR (Table S2). Separate PCR reactions were conducted for each sample, while gDNA and eDNA samples were processed in separate experiments, with negative controls (PCR blanks) included for each plate. Multiplex amplicon PCR products were assessed for successful amplification and size on TapeStation D1000 before proceeding. All amplicon products were diluted to 1:10 and indexed with Illumina DNA/RNA Unique Dual (UD) Indexes during a second-stage PCR (Table S1; Table S3). Indexed sample concentrations were determined using the Qubit dsDNA HS (High Sensitivity) kit on a Qubit 4 fluorometer, before samples were pooled at equimolar concentrations. A bead cleanup was then performed on the pool with Agencourt AMPure XP beads at 0.8 x pool volume to remove small fragments. The clean pooled library was sequenced on a 25M 600 cycle Illumina MiSeq i100 Series cartridge with 5% PhiX spike in, targeting 1000 reads per marker per sample.

#### 2.2.4 Bioinformatics workflow

Sequence reads for each sample were demultiplexed with the MiSeq Analysis Software (Illumina) and run through fastp to remove any remaining Illumina adapters and low-quality reads (Chen et al., 2018). PCR primers were removed with cutadapt (Martin, 2011), before read pairs were merged with fastp (minimum overlap 100 base pairs). BWA-MEM2 (Vasimuddin et al., 2019) was used to map merged reads to a reference fasta of the targeted amplicon sequences derived from the reference assembly. The resulting BAM files were sorted and indexed with samtools (Danecek et al., 2021). Samples where no reads successfully mapped to a reference microhaplotype sequence were removed from further analysis. SNPs were called separately for all remaining tissue and eDNA derived samples with BCFtools (Danecek et al., 2021) *mpileup* and *call* (--skip-variants indels-mv) functions. Called SNPs were filtered for a Hardy-Weinberg equilibrium of 0.001 using VCFtools (Danecek et al., 2011) to remove likely sequencing errors and a minimum mean SNP depth of 10 with BCFtools (-e ‘MEAN(FMT/DP) < 10’). Microhaplotype markers with no SNPs that passed filtering were removed from further analysis Microhapulator (Standage et al., 2022) was then run with the final bam files and SNP dataset to generate microhaplotypes for each marker per sample. This output was converted into two matrices containing read counts of each allele across all tissue and eDNA samples separately. Any alleles with read count under 5 were discarded and then read counts were normalised within each sample to account for difference in sequencing depth before analysis.

### 2.3 Efficacy of microhaplotype markers for inferring population genetics parameters from target eDNA samples

To begin investigating the possible use of microhaplotypes for population genetic inferences, we aggregated normalised read counts across all tissue samples for each allele to create population-level allele frequencies. This process was then repeated for the target eDNA samples separately. Population allele frequencies for each microhaplotype were calculated for both tissue and target eDNA-derived samples and evaluated for correlation using Pearson’s product-moment correlation and Spearman’s rank correlation statistics with the *cor.test* function in the R stats v4.5.1 package (R Core (Team, 2024). This test was repeated for allelic richness, calculated as the number of unique haplotypes per marker across all tissue and target eDNA samples. We also calculated microhaplotype allele frequencies for each individual sample to assess the concordance of genetic profiles derived from tissue and corresponding eDNA samples. Allele frequencies under 1% were removed to reduce impacts of sequencing errors. Similarity was visualised between allele frequency profiles from eDNA and corresponding individual tissue samples using a dendrogram and Principal Components Analysis (PCA). The dendrogram was based on Euclidean genetic distances, generated using *dist* and hierarchically clustered, while the PCA was performed with the *prcomp* function, both in R stats. Genetic profile concordance between eDNA and tissue-derived samples was assessed both with replicate samples calculated separately and with replicates pooled per site for eDNA samples.

### 2.4 Sensitivity analyses

The discriminatory power of the microhaplotype panel was evaluated by calculating the probability of identity (*P_ID_ = sum_i p_i^4^ + sum_(i<j) (2 p_i p_j)^2^*) and probability of identity among siblings (*PI_sibs_ = 0.25 + 0.5 x sum_i p_i^2^ + 0.5 x (sum_i p_i^2^)^2^ - 0.25 x sum_i p_i^4^*) for each marker using population allele-frequencies derived from the 21 tissue samples (Taberlet & Luikart, 1999). Cumulative *P_ID_* and *PI_sibs_* were then calculated across the marker panel, representing the probability that two unrelated individuals or two siblings, respectively, would share identical multilocus genotypes across all markers included up to each successive marker. These cumulative estimates were used to assess the overall discriminatory power of the microhaplotype panel.

To assess the minimum number of microhaplotype markers needed for reliable assignment of eDNA samples to their corresponding tissue-derived reference samples, we compared allele-frequency profiles across all markers shared between eDNA and tissue samples. For each microhaplotype marker, similarity between an eDNA sample and each candidate tissue sample was calculated as the sum of the minimum allele frequencies across all alleles at that marker, with values ranging from 0 (no shared allele-frequency profile) to 1 (identical allele-frequency profiles). Similarity scores were then averaged across the selected markers, giving equal weight to each marker. The tissue sample with the highest mean similarity score was considered the predicted match. Assignment accuracy was then calculated as the proportion of eDNA samples for which the known corresponding tissue sample was ranked as the highest-scoring candidate. To evaluate the effect of panel size on individual assignments, random subsets of 1 to the maximum microhaplotype markers were selected, with 100 independent random subsets evaluated for each marker number. For each random marker subset, all eDNA samples were compared against all available tissue samples. Mean assignment accuracy and the 2.5th and 97.5th percentiles across random marker subsets were calculated for each marker number. We identified the minimum marker thresholds required to achieve 95% and 99% mean assignment accuracy as the smallest number of markers for which mean assignment accuracy reached or exceeded the respective threshold. A conservative marker threshold was also calculated as the minimum number of markers for which the lower 2.5th percentile, or lower bound, of the empirical distribution of assignment accuracy reached the 95% and 99% thresholds.

Similarity margins were calculated for each eDNA sample as the difference between the similarity score of the known corresponding tissue sample and that of the second highest-scoring tissue sample. Thus, larger positive margins indicated greater separation between the true tissue match and the nearest alternative, whereas margins close to zero indicated less distinct assignments. Mean similarity margins were calculated across eDNA samples for each random marker subset and used as a complementary measure of discriminatory power. We calculated the 2.5th and 97.5th percentiles of mean similarity margins across random marker subsets for each marker number to characterise the distribution of discriminatory performance.

### 2.5 Non-target eDNA assignment

For each non-target eDNA sample, we followed the same process to create a dendrogram and heatmap based on Euclidean distance to assess whether samples clustered with any known tissue reference individuals. To assess assignment stability, 100 bootstraps of random subsets from 1 to the maximum number of recovered markers were evaluated for each sample, with marker subsets restricted to those recovered in that sample. For each bootstrap, we calculated similarity to all available tissue reference profiles using the frequency-overlap metric, defined as the sum of the minimum allele frequencies between the eDNA and tissue profiles, averaged across markers, and ranked candidates by similarity. Assignment stability was calculated as the proportion of replicates in which each reference individual was the top candidate, whereas we assessed assignment margin from the mean difference in similarity between the top two candidates. Final assignments for each unknown eDNA sample were based on all markers recovered, with the two highest-ranking reference individuals and the difference in their similarity scores reported. Only samples with high similarity and clear differentiation between the top two candidates were considered as successful matches.

### 2.6 Comparative analysis of metrics derived from tissue and eDNA samples

Population genetics metrics derived from microhaplotype data from tissue and targeted eDNA samples were compared with those derived the WGS data for the same 21 individuals to test whether microhaplotype markers produced comparable genome-wide inferences. Resequencing data was aligned to the tiger shark reference genome assembly (Pearce, Doran, et al., 2026) with bwa-mem2 v2.2.1 (Vasimuddin et al., 2019). Duplicate reads were removed with Picard’s *MarkDuplicates* function within the GATK v 4.2.5.0 package (McKenna et al., 2010). BCFtools v1.15 (Danecek et al., 2021) *mpileup* and *call* functions were then used to call SNP variants across these samples. Only biallelic SNPs with a minimum mean 10X depth were retained after filtering using VCFtools v0.1.16 (Danecek et al., 2011). Given the limited sample size (n = 21), we filtered variants using a minor allele count (MAC) threshold of 3 to remove singletons and doubletons and reduce the influence of poorly estimated rare alleles, rather than a fixed minor allele frequency threshold. This pipeline was repeated for the tissue and eDNA-derived microhaplotype dataset but aligned to a reference microhaplotype library. To then account for linkage disequilibrium (LD), three LD filtering schemes were applied across all datasets: baseline (no LD filter), distance-based thinning (VCFtools; WGS: 10kb; MH: 250bp), and pruning (r^2^=0.2). LD pruning was conducted using the SNPRelate v1.32 R package (Zheng et al., 2012) with an r^2^ threshold of 0.2. DartR (Gruber et al., 2018; Mijangos et al., 2022) *gl.report.heterozygosity* was used to calculate global unbiased expected heterozygosity (uHe) and Inbreeding coefficient (FIS), as well as individual-level observed heterozygosity (Ho) for the three filtering schemes of each dataset. Pearson’s correlations between the same individuals across all datasets and filtering schemes were then assessed. Principal component analysis (PCA) was performed using the *snpgdsPCA* function from SNPRelate using the pruned datasets only according to the package recommendations.

### 2.7 Validation datasets

#### 2.7.1 Acoustic telemetry

All tiger sharks sampled in this study were identified with internal acoustic tags that reported to receiver arrays within Exmouth Gulf (Gascoyne) and the Mid-West regions of Western Australia. In Exmouth Gulf, sharks were tagged in October 2023 and acoustic data from the receiver array was retrieved 12 months later in October 2024. Additional sharks were tagged in this region in 2025; however, no further acoustic data was available at time of analysis. Tiger sharks from the Mid-West region (Cervantes, Jurien Bay and Geraldton) were tagged in May-July 2024, with acoustic data downloaded from Cervantes and Jurien Bay stations in October 2025 and Geraldton in April 2026. We used the raw acoustic detections to validate any non-targeted eDNA microhaplotype results.

#### 2.7.2 eDNA metabarcoding

Metabarcoding data were available only for samples collected in 2024 within Exmouth Gulf in the Gascoyne region (*n sites* = 6, *n samples* = 24). We processed the same samples through both the microhaplotype panel and standardised metabarcoding workflows after initial DNA extraction (*2.2.2 Sample laboratory processing*). The metabarcoding workflow adhered to the standardised and automated OceanOmics Centre marine vertebrates protocol (Missen, Huynh, et al., 2026), applying three metabarcoding assays (16S-Fish/D, MiFish-U/E2 (multiplexed), and MarVer1). These samples were sequenced on a NextSeq 2000. A publicly available, automated Nextflow bioinformatic pipeline (Pearce et al., 2023) (https://github.com/Minderoo-OceanOmics-Centre-UWA/OceanOmics-amplicon-nf) was then implemented to produce the final taxonomically annotated Amplicon Sequence Variants (ASVs). For this study, we retained only tiger shark taxonomic assignments and compared them with acoustic telemetry and microhaplotype datasets.

## 3. RESULTS

In total, paired tissue and targeted eDNA samples were collected from 21 individual tiger sharks between 2023 and 2025 from the Gascoyne and Mid-West regions of Western Australia (Figure 1). We also collected non-targeted eDNA samples opportunistically when no tiger sharks were caught but they were likely to be in the vicinity, to test the feasibility of microhaplotype recovery from in situ aquatic samples. This resulted in a total of 84 targeted eDNA samples (four replicates per individual) and an additional 43 non-targeted eDNA samples (11 sites).

### 3.1 Individual identification from targeted eDNA

MiSeq 300bp paired-end sequencing produced similar microhaplotype recovery (mapped reads and total marker number) between paired tissue and targeted eDNA samples (Table S4). Seven out of the 55 microhaplotype markers did not contain any expected variants. Filtering for HWE and 10X mean depth reduced this to a set of 42 markers for microhaplotype calling. No laboratory controls (extraction and PCR blanks) or field controls from Exmouth Gulf in 2024 when no sharks were tagged retained reads after mapping to the reference microhaplotype amplicon library. Four markers did not produce sufficient depth across samples to reliably call microhaplotypes, resulting in a final set of 38 markers. Both tissue and targeted eDNA samples averaged 36 markers per sample following microhaplotype calling and depth filtering. However, field controls from trips which involved the capture of sharks displayed low levels of contamination throughout.

The final microhaplotype panel contained 151 observed allele combinations (phased microhaplotypes), with near-perfect correlations between paired tissue and eDNA samples. Microhaplotype allele frequencies (AF) calculated across all samples from tissue and targeted eDNA were strongly and significantly correlated (Figure 2.A; Pearson’s product-moment correlation: 0.997, p-value < 0.001; Spearman’s rank correlation rho: 0.993, p-value < 0.001). Markers ranged from one to six SNPs, which formed up to 8 (eDNA; tissue = 7) microhaplotype allele combinations per marker (allelic richness; AR) after filtering out potential sequencing errors (AF > 0.01). Microhaplotype allelic richness was steadily inflated in targeted eDNA samples relative to tissue, even when frequencies below 1% were filtered out (Figure 2.B). Regardless, there remained a significant strong correlation, as this bias was concentrated at highly polymorphic markers (Figure 2.B; Pearson’s product-moment correlation: 0.98, p-value < 0.001; Spearman’s rank correlation rho: 0.98, p-value < 0.001).

**Figure 2.**
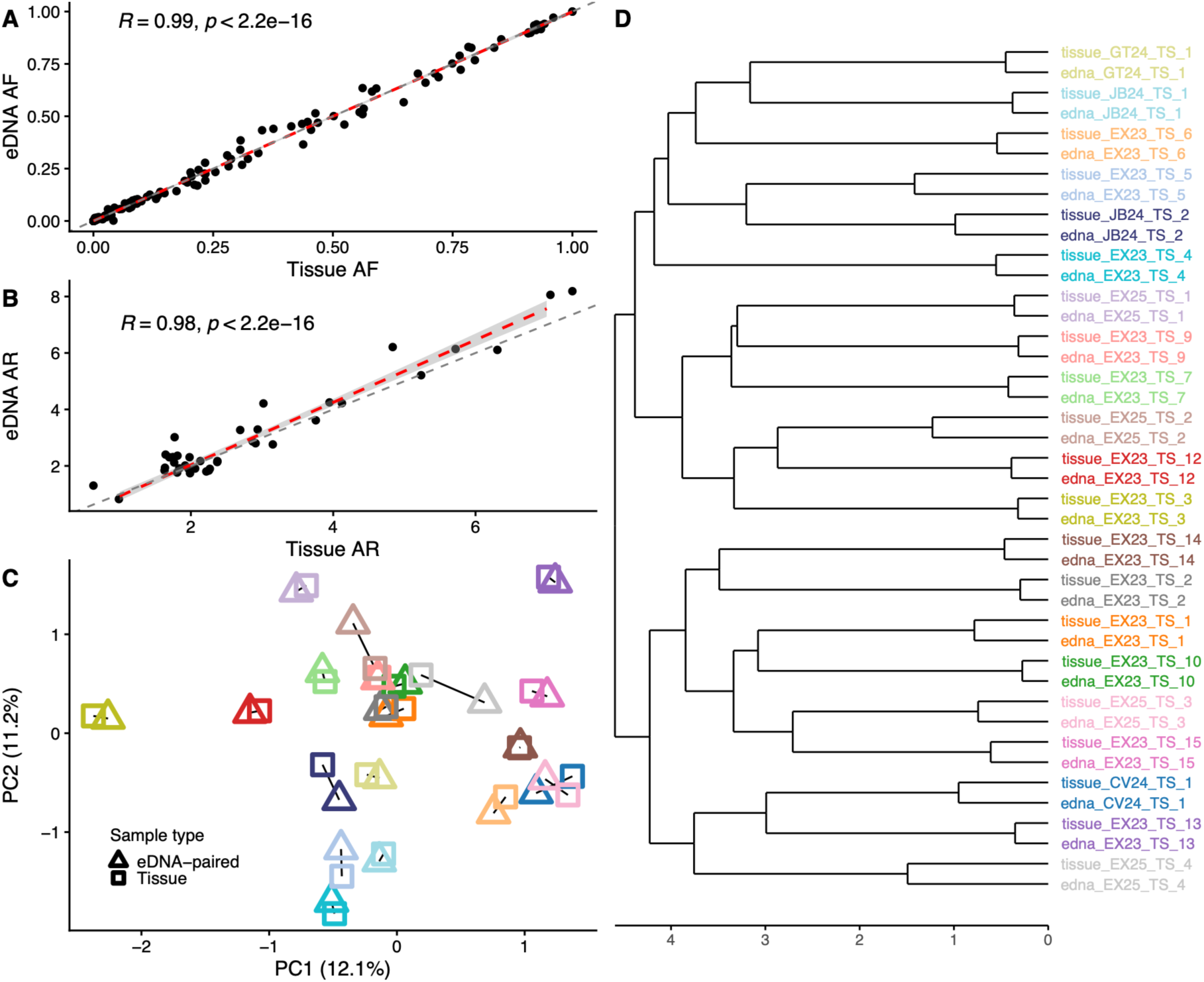
Tissue and eDNA-derived tiger shark microhaplotypes: A) Allele frequencies (AF) calculated across all samples for each marker along with the Spearman’s rank correlation coefficient B) Total allelic richness (AR) per marker between sample types and the Spearman’s rho, C) Principal Component Analysis (PCA) plot colour-coded by individual sharks, with solid black lines connecting sample pairs that correspond to the same individual and D) dendrogram constructed using Euclidean genetic distances displaying the clustering between paired samples from the same individual shark. Sample colours in the PCA correspond to those in the dendrogram.

Furthermore, microhaplotypes could be matched reliably between tissue-derived and targeted eDNA water samples taken close to the same individual tiger shark. The PCA plot showed that paired tissue and eDNA samples from the same individual generally clustered closely together (Figure 2.C). A dendrogram constructed using Euclidean genetic distances showed all associated samples paired together (Figure 2.D). The majority of eDNA replicates remained clustered with their corresponding tissue sample within the dendrogram and PCA when allele frequencies were calculated for each replicate separately (Figure S1; Figure S2). However, the individuals EX25_TS_2 (tan), EX25_TS_4 (grey) and JB24_TS_2 (navy blue), displayed greater separation between tissue and corresponding eDNA samples within the PCA, both as pooled and individual replicates (Figure 2.C; Figure S2). Individual EX25_TS_3 was the only exception that did not fully cluster in the dendrogram when calculating eDNA replicates separately (Figure S1). This shark showed large replicate variation in the PCA (Figure S2) that was resolved when replicates were pooled. Thus, to ensure robustness, we pooled eDNA replicate allele frequencies for all analyses.

### 3.2 Sensitivity analyses

The phased microhaplotypes from tissue samples provided high discriminatory power for individual identification. The cumulative probability of identity (P_ID_) for unrelated individuals was 4.90 × 10⁻¹⁴, corresponding to approximately one expected match among 20 trillion (2.0 × 10¹³) unrelated individuals. The more conservative cumulative probability of identity among full siblings (PI_sibs_) was 4.68 × 10⁻⁷, corresponding to approximately one expected match among 2.1 million sibling pairs.

To evaluate the minimum number of microhaplotype markers required to achieve accurate results, we calculated the average assignment accuracy (proportion of correctly identified samples) across 100 bootstraps of random marker subsets from one to 38 microhaplotype markers. Assignment accuracy increased with the number of microhaplotype markers included, reaching an average of 95% of samples correctly identified with just nine markers (Figure 3. A), although 13 markers were required for the lower 2.5th percentile (i.e. lower 95% bound; conservative threshold) to reach this threshold (Figure 3. A). Mean assignment accuracy exceeded the 99% threshold with 19 microhaplotype markers, but 27 markers were needed to reach the conservative threshold (Figure 3. A). Assignment confidence was also assessed by comparing how similar the top two candidate tissue samples were for each eDNA sample assignment. The margin between the known tissue match and the second-best candidate increased with microhaplotype marker number (Figure 3. B). This reiterated that increasing marker numbers improved assignment accuracy, confidence and discriminatory power for sample identification.

**Figure 3.**
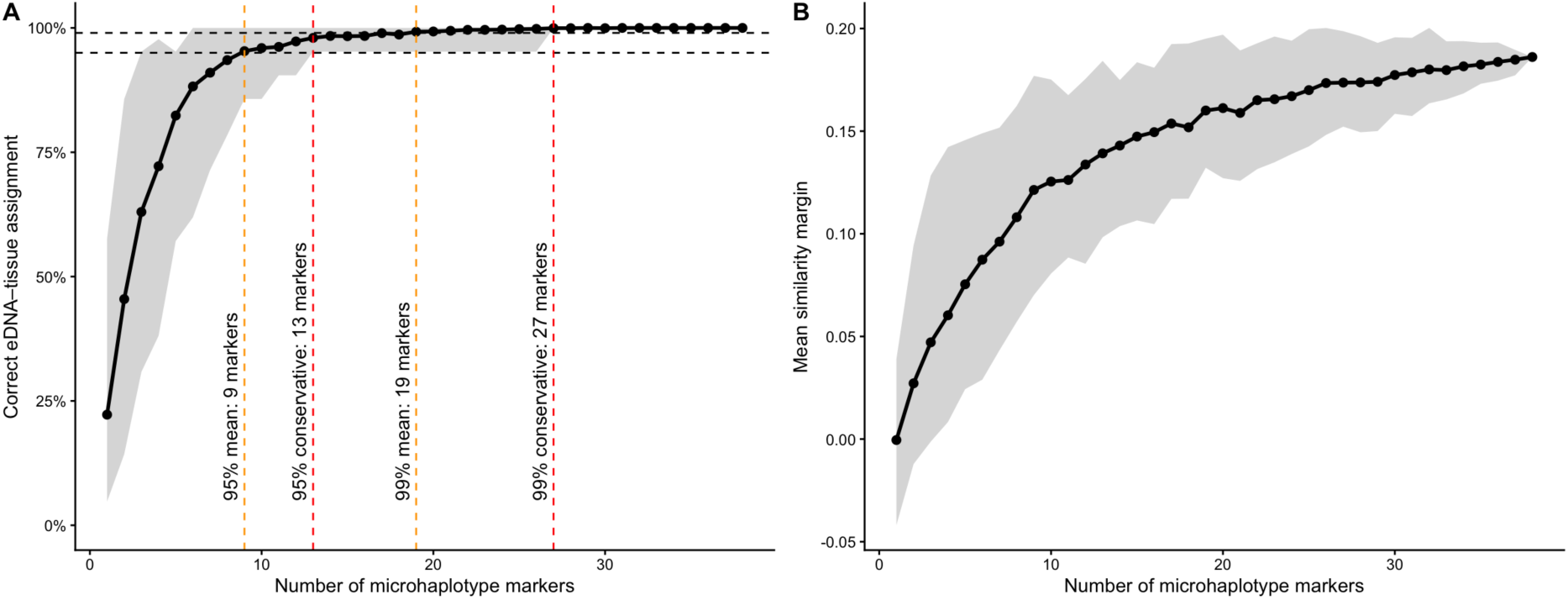
Microhaplotype marker thresholds required to correctly match allele frequencies from eDNA and tissue samples. A) Proportion of eDNA samples correctly identified out of 100 bootstraps. Thresholds were set when the mean assignment accuracy (orange) and the lower quantile (conservative threshold = red) reached 95% and 99% of samples correctly identified. B) Separation between true and alternative tissue matches with increasing microhaplotype marker number. Calculated as the margin between the mean allele frequency similarity of the true match and the second-best tissue sample selected.

### 3.3 Population genetics metrics from microhaplotypes

Population-level estimates were calculated from both microhaplotype sample types and compared to estimates derived from whole-genome sequencing (WGS) data for the same individuals to ascertain the panel’s use for population genetics applications. Principal Components Analysis (PCA) revealed no clear clustering by region or sampling year evident in any dataset, with the microhaplotype-derived PCAs broadly recapitulating the lack of genetic structure observed in the WGS data (Figure 4.B). Therefore, we assumed all samples belonged to a single population. Additionally, due to the nature of microhaplotypes, linkage disequilibrium (LD) effects were assessed through three filtering schemes: baseline (no LD filter), distance-based thinning and r^2^ pruning (Table S5). LD was only considered within microhaplotype markers, as the majority of markers were located on different chromosomes. Chromosomes 3, 5, 8, 18 and 37 each contained 2 markers (distance ranged 522.23kb – 64.43Mb), however chromosomes 31 and 34 had 5 markers each, but all were a minimum of 60kbp apart (Chr31 distance range: 67.9kb – 17.55Mb; Chr34: 147.5kb – 25.04Mb). Pruning retained 39 SNPs for both microhaplotype datasets, whereas thinning kept one per marker which resulted in 36-37 SNPs retained per sample.

**Figure 4.**
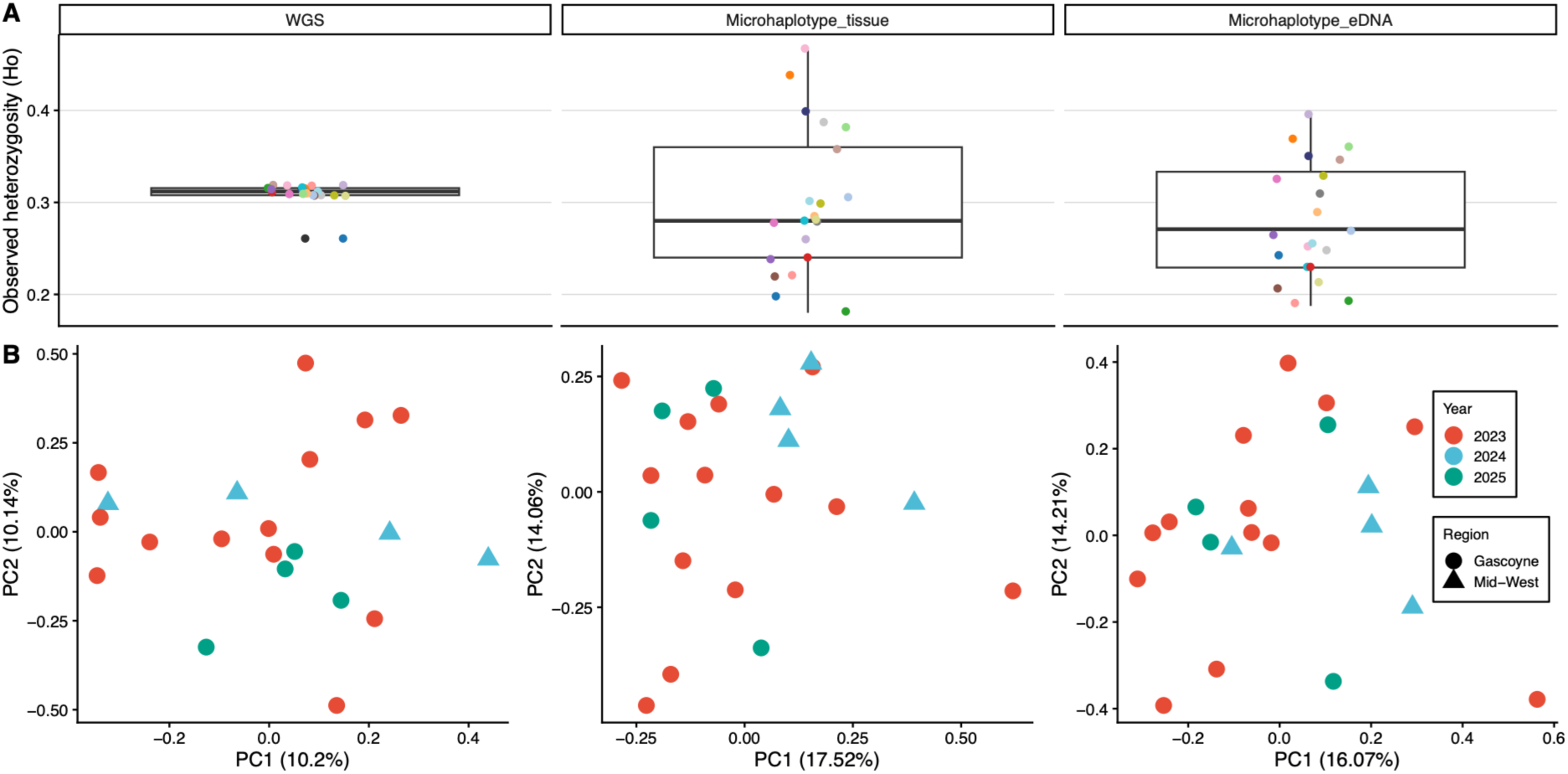
Population estimates of tiger shark samples derived from 10X whole-genome sequencing (WGS), tissue-based and eDNA water sample microhaplotyping. A) Observed heterozygosity (Ho) per individual, B) Principal Components Analysis (PCA) of individuals across regions and years from WGS, microhaplotype tissue samples and targeted eDNA samples (left to right). Linkage disequilibrium (LD) pruning (r^2^=0.2) was applied across all datasets.

Estimates of unbiased expected heterozygosity (uHe) were broadly consistent across the three datasets, irrespective of SNP filtering (Table S6; uHe = 0.272 - 0.341). Inbreeding coefficients (FIS) based on WGS data remained close to zero across all filtering schemes (Table S6). FIS estimates from tissue-derived microhaplotypes dropped after pruning but not thinning, suggesting the panel provides comparable, but potentially inflated, estimates of population genetic diversity when accounting for linkage among SNPs. However, eDNA-derived microhaplotypes showed persistently elevated FIS that pruning or thinning did not resolve (0.144–0.210), indicating a cause independent of LD.

Observed heterozygosity (Ho) per individual differed substantially between WGS and microhaplotype data (Figure 4.A). WGS-derived Ho was very consistent across individuals, reflecting the greater number of loci. In contrast, both eDNA and tissue microhaplotypes showed considerable variance in per-individual Ho (Figure 4; tissue: 0.162 - 0.538; eDNA: 0.128 - 0.417), consistent with the small number of loci and higher sampling variance inherent to panel-based genotyping. This pattern was reflected across all three filtering approaches (Figure S3). At an individual level, Ho was significantly correlated for the same samples between microhaplotype datasets across all filtering schemes, but this relationship strengthened when LD was accounted for (Figure S4–S6: r = 0.499 - 0.627; p < 0.01). Conversely, Ho displayed consistently weak, non-significant correlations between WGS and microhaplotype datasets, regardless of filtering scheme (Figure S4–S6: r = 0.114 - 0.304). This indicated that while tissue and eDNA microhaplotypes capture concordant individual heterozygosity estimates, microhaplotype-derived Ho does not reliably reflect genome-wide individual heterozygosity as estimated from WGS (Figure S4–S6).

### 3.5 Non-targeted eDNA sampling

To test the validity of this technique for broadscale monitoring applications, 43 eDNA samples were also taken from 11 sites without the visual confirmation of tiger shark presence. As expected, these non-targeted eDNA samples showed higher variability in reads and marker recovery (Table S4). Metabarcoding sequence data were only available for six of these sites, all containing positive detections of tiger sharks (Table 1). Five sites had no microhaplotype markers after depth filtering, despite a positive acoustic detection at one of those sites (Table 1). Microhaplotypes were successfully recovered from 21 separate samples across six non-targeted sites (Table 1). Three sites (EX23_BS, CV24_CV and GT24_GTS) produced a confident match to a tiger shark from our reference tissue dataset (Figure 5). When compared against all potential reference individuals, these samples displayed high candidate similarity (Table S7) and were reasonably distinct from the next top candidate (Table S7; Figure S8). Assignment stability also plateaued when replicated over 100 bootstraps of varying marker subsets, indicating confidence and sufficient marker recovery (Figure S7). Furthermore, we validated two of these eDNA assignments with a positive ID match to an acoustic tag detection at the corresponding location that day, and the third with a shark tagged nearby the same day (Table 1).

**Figure 5.**
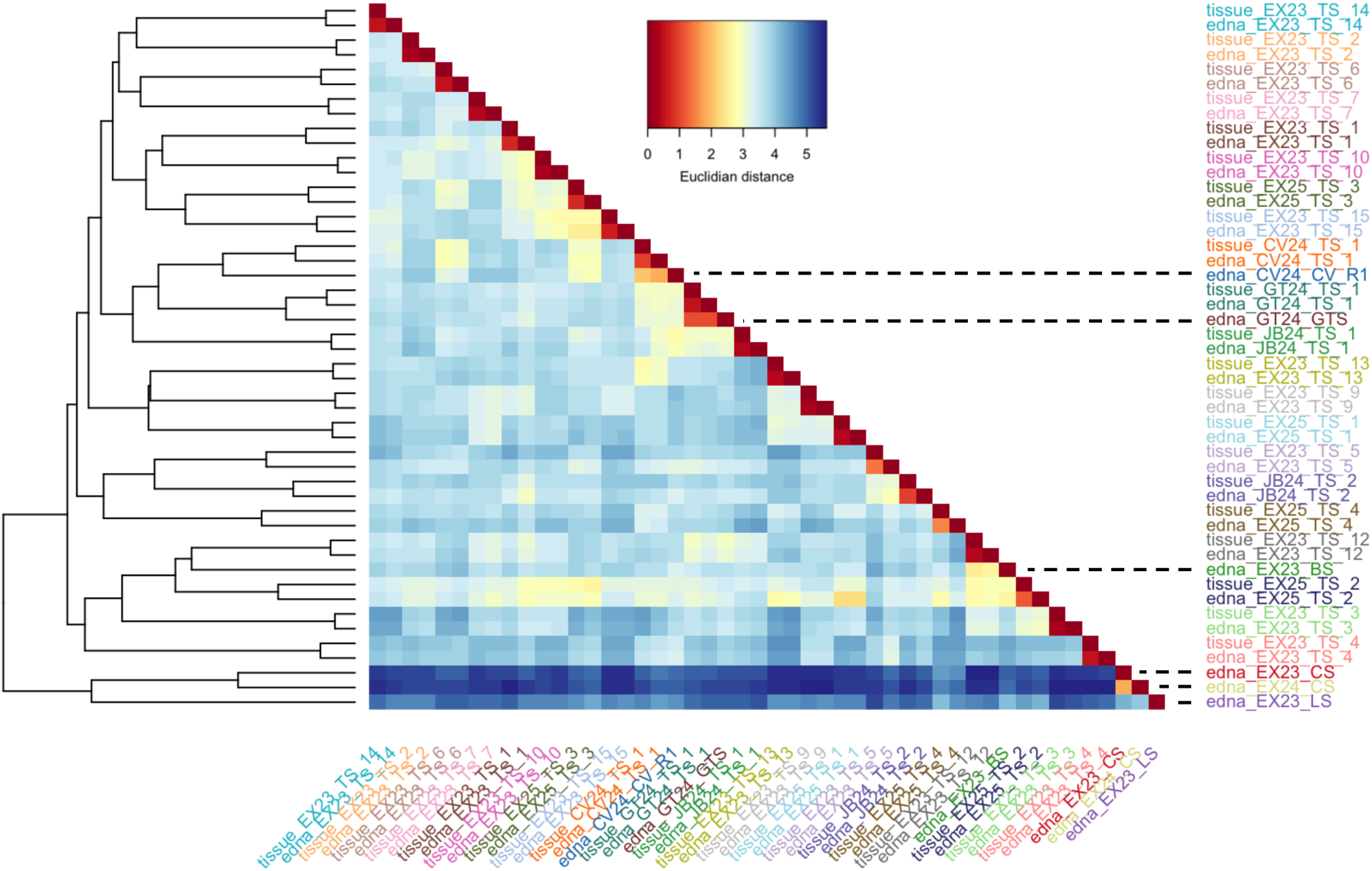
Comparison of non-targeted ‘background’ eDNA samples taken from known tiger shark areas but without presence records. Samples are colour-coded by sampling site/individual shark, with pooled eDNA sample replicates (*n* = 4). The dendrogram (left) shows sample clustering, whereas the heatmap (middle) shows pairwise similarity. Black dotted lines highlight where the six background eDNA sampling sites sit.

**Table 1.** Tiger shark detections within non-targeted eDNA water samples across three different methods: nuclear microhaplotyping (MH) after filtering, mitochondrial metabarcoding (MB) and acoustic tags. N. samples refers to the number of replicate samples with positive detections for that site (*max N* = 4). N. markers refers to the number of markers successfully microhaplotyped for that site across all replicates. Acoustic detections were only included for the same day water sampling was conducted. *N/A* = Not Available. Detection times are in 24-hour format.

| **Site** | **Number of MH** |  | **N. MB samples** | **Acoustic detection** |  | **eDNA assignment** |  |
|---|---|---|---|---|---|---|---|
|  | Samples | Markers |  | ID | Time | ID | Time |
| **EX23_BS** | 4 | 31 | *N/A* | EX23_TS_12 | 16:00-17:00 | EX23_TS_12 | 16:00 |
| **EX23_CS** | 3 | 6 | *N/A* | No | - | No | 16:00 |
| **EX23_LS** | 4 | 17 | *N/A* | EX23_TS_13<br>EX23_TS_3 | 14:30-16:30<br>14:30 | No | 16:00 |
| **EX24_MB** | 0 | 0 | 2 | No | - | No | - |
| **EX24_BS** | 0 | 0 | 3 | EX23_TS_7 | 13:00 | No | 14:00 |
| **EX24_VLF** | 0 | 0 | 2 | No | - | No | - |
| **EX24_CS** | 2 | 2 | 3 | No | - | No | - |
| **EX24_BOR** | 0 | 0 | 1 | No | - | No | - |
| **EX24_SS** | 0 | 0 | 1 | No | - | No | - |
| **CV24_CV** | 4 | 33 | *N/A* | No | - | CV24_TS_1 | 15:00 |
| **GT24_GTS** | 4 | 37 | *N/A* | GT24_TS_1 | 17:00 | GT24_TS_1 | 17:30 |

We recovered too few microhaplotype markers from two sites where there were no acoustic detections to make an identification (Table 1). EX23_CS lacked metabarcoding data to corroborate this, but positive detections of tiger sharks were found at EX24_CS. Markers from these sites could not be assigned to a known individual as the microhaplotypes did not cluster (Figure 5) and had inconsistent assignment stability that did not reach a plateau due to exceedingly small marker recovery and margins between candidate reference individuals (Table S7; Figure S8). Site EX23_LS also still could not be assigned to a reference despite higher microhaplotype recovery and acoustic confirmation of known individuals (Table 1). Interestingly, the top candidate assignment for this site (EX23_TS_9) was not one of these acoustic detections, whereas the second-top candidate assignment was EX23_TS_13. However, there was minimal distinction (0.01) between candidates (Table S7; Figure S8),) and assignment stability did not plateau sufficiently (Figure S7) to confidently assign a candidate.

## 4. DISCUSSION

Monitoring shark populations in Western Australia and worldwide currently relies on conventional methods that are resource-intensive and limited in spatial and temporal scale (Carrier et al., 2018; D’Antonio et al., 2025; Heithaus et al., 2007). With eDNA emerging as a non-invasive, potentially scalable, cost-effective alternative, this study examined whether a microhaplotype panel specific to tiger sharks could serve as a versatile genomic tool for both population genetic inferences and individual identification from conventional and environmental samples. The final 38-marker panel, comprising 151 variant sites, enabled highly accurate individual discrimination from paired tissue and eDNA water samples and broadly recapitulated population genetic patterns identified using WGS. Application to non-targeted seawater samples further demonstrated the feasibility of recovering individual-specific genetic information from in-situ eDNA, with three samples assigned to known individuals, two of which were independently supported by concurrent acoustic detections. However, variable marker recovery from seawater, along with possible mixed contributors, prevented individual identification in other samples. These results underscore the constraints of applying nuclear DNA techniques to environmental samples and the challenges for extending this approach to broader monitoring applications. Nevertheless, these findings show that microhaplotypes can bridge population genetics and individual-level monitoring, offering a promising approach to integrate genomic identification and enable more scalable, less invasive monitoring of wildlife populations.

Our results provide a proof-of-concept for individual-level identification of megafauna from non-targeted eDNA samples, demonstrating the potential to extend environmental DNA monitoring beyond species detection. Of 11 non-targeted sampling sites, microhaplotypes were recovered at six, with three samples confidently matching known individuals. At both Mid-West sites (Cervantes and Geraldton), eDNA matched the same individual that had been tagged earlier that day, despite water samples being collected at least 4 km from the tagging locations. However, the Cervantes shark was tagged 10 km away from the receiver and so was not acoustically detected. Similarly, at Bennet Shoal in 2023, eDNA matched an individual tagged six hours prior and 15 km away at Larkin Shoal, with acoustic detections independently confirming its presence at the time of sampling. For all non-target eDNA samples, fresh gloves, and sterilised collection bottles and filtration cups that had not been previously used that day were employed to minimise potential contamination. These assignments showed high candidate similarity and clear margins between competing individuals, providing compelling evidence in the identifications. Although false positives are also an important consideration in eDNA-based approaches (Goldberg et al., 2016), if this were the cause we would expect these to occur more frequently in non-target samples collected on tagging days, which was not observed. Importantly, although mitochondrial metabarcoding detected tiger shark DNA at all non-targeted sites, most yielded no nuclear microhaplotypes or concurrent acoustic detections. This suggests that mitochondrial DNA may remain detectable longer than the nuclear fragments targeted by the microhaplotype assay, resulting in a narrower temporal and spatial window for individual identification than for species detection. This presents a challenge for scaling this approach beyond this study before it can be reliably applied to broader monitoring programmes.

We found that microhaplotypes provided efficient individual-level genetic identification in tiger sharks, derived from both conventional tissue and targeted environmental samples. The overall strong concordance in allelic frequencies, richness and individual assignments between paired tissue and eDNA samples reflects that found in previous microhaplotype (Meenakshisundaram, Jarman, et al., 2025; Meenakshisundaram, Meekan, et al., 2025) and other environmental studies (Barba et al., 2023). Three sets of paired samples notably deviated within both pooled and replicate PCAs, but still clustered together within both dendrograms (Figure 2). Individual EX25_TS_3 was the singular exception where a few replicate samples did not cluster with the corresponding tissue sample (Figure S1), reflecting the large variation seen in the PCA (Figure S2). However, pooling eDNA replicates resolved this issue. A small fraction of loci differed in allele frequencies, driving the separation despite the majority matching precisely, likely due to low-level contamination seen in the field controls or multiple contributors. However, any contamination that was present in these samples was not severe enough to impact the outcomes as overall genetic profile still matched between tissue and targeted eDNA sample pairs. Successful results have still been demonstrated from a singular eDNA sample (Meenakshisundaram, Meekan, et al., 2025) and although our findings largely agree, this emphasises that pooling eDNA replicate allele frequencies increases robustness when dealing with an inherently highly variable sample type. Overall, this attests that genetic information comparable to that obtained from conventional tissue samples can be recovered from environmental samples, offering an alternative to invasive sampling and a potentially method for monitoring in the absence of visual census.

Our microhaplotype panel demonstrated high discriminatory power for individual identification across unrelated individuals and among full siblings, consistent with previous evaluations (Baetscher et al., 2018; Tomas et al., 2024). Assignment correctness (accuracy) and margin between top two candidates (confidence) both increased with the number of microhaplotype markers (Figure 3). This reiterates that sufficient marker recovery is not only essential for robust individual discrimination, but also to reduce assignment ambiguity when candidate reference profiles are only marginally different. Sensitivity analyses indicated that at least 13 markers were required to correctly identify 95% of individuals under the more conservative threshold, increasing to 27 markers to achieve 99% accuracy. Yet, Larkin Shoal yielded 17 markers, and two reference-genotyped sharks were acoustically present at the site, but lacked sufficient differentiation between candidate individuals. This may instead reflect contributions from multiple individuals, an individual absent from the reference dataset or insufficient genetic information for confident assignment. Furthermore, the relatively narrow and overlapping similarity margins observed in other samples between true matches and second candidates indicates that assignment confidence cannot be determined solely from the highest similarity score. Although we recommend 27 markers as a preliminary conservative threshold, these estimates were derived from a relatively small reference dataset and discriminatory power and assignment confidence may change with larger reference populations, unknown individuals or varying mixed samples, particularly where closely related individuals are included. Still, this demonstrates that relatively small, targeted panels can provide sufficient resolution.

The concordance between microhaplotype and WGS derived population genetics metrics suggests that the panel captures broad patterns of genetic variation, in this case reflecting the high connectivity and lack of population structure characteristic of highly mobile marine predator (Bernard et al., 2021; Pirog et al., 2019). However, elevated FIS in eDNA samples persisted after accounting for LD, suggesting that this pattern reflects factors such as uneven allelic representation and allelic dropout from low-template DNA (Andres et al., 2021; Brandão-Dias et al., 2026; Couton et al., 2023). The reduction in FIS following LD pruning in tissue samples also highlights the importance of accounting for linkage among SNPs within microhaplotype markers (Couton et al., 2023). Microhaplotype-derived individual observed heterozygosity was more variable than WGS estimates because it is estimated from fewer loci, which increases sampling variance (Couton et al., 2023). Nevertheless, the strong correlation between matched tissue and eDNA estimates indicates that the panel provides consistent individual-level estimates of conventional and environmental microhaplotypes but should not be considered a proxy for genome-wide heterozygosity. Overall, the panel reliably captured population-level genetic diversity from both tissue and targeted eDNA, while individual-level estimates are more sensitive to the reduced genomic representation of the panel and characteristics of eDNA sampling. This suggests that microhaplotypes could be a cost-effective alternative for population genetics from conventional tissue and targeted environmental samples. However, retaining a single SNP per marker limits population genetic analyses to biallelic, frequency-based metrics, whereas incorporating multi-allelic microhaplotype genotypes could provide greater discriminatory power and additional resolution of genetic variation.

Insufficient genetic variation to construct microhaplotypes from 17 of the 55 sequenced markers likely reflects a combination of low sequencing depth and limited genetic diversity among the sampled tiger sharks, given that the panel was designed using individuals spanning the broader Indian Ocean. Consequently, the panel may capture greater variation across more geographically distant populations, highlighting the importance of considering population structure and diversity during panel design (Meenakshisundaram, Jarman, et al., 2025; Solari et al., 2024). Given the pronounced genetic distance between the Indo-Pacific and Atlantic tiger shark metapopulations (Holmes et al., 2017; Pirog et al., 2019), the current panel is likely most representative of Indo-Pacific populations and may require further optimisation for Atlantic applications. Of the initial 80 markers developed, 55 passed laboratory optimisation and 38 were retained for analysis, emphasising further optimisation is required and the value of incorporating redundancy into future panel designs. Furthermore, the development of multi-allelic microhaplotype genotype analysis would provide greater resolution of within-population and individual-level genetic variation for population genetic analyses over bi-allelic SNP frequency-based metrics for future applications. The use of inexpensive, standard amplicon sequencing nevertheless makes these panels readily scalable for cost-effective population and individual genetic monitoring.

As environmental DNA approaches advance, remaining technical challenges increasingly reflect biological constraints, particularly the persistence and availability of nuclear DNA following shedding (Jo & Minamoto, 2021; Jo et al., 2022). Our results highlight this challenge, although filtering larger volumes of water may increase DNA recovery when target animals are present, this relationship appears context-dependent (Barba et al., 2023), and our limited results suggest that successful individual identification may require sampling within a relatively short period following animal presence within an aquatic environment. Mixed-individual samples and contamination present an additional challenge, as demonstrated at Larkin Shoal, and are likely amplified in non-target eDNA samples, where low template concentrations increase these effects. Scaling the approach will therefore require future validation to assess assignment performance using larger, geographically representative reference datasets containing related and unrepresented individuals, and establish confidence thresholds as well as species and environment-specific detection windows. Single use filtration equipment or at least further optimised sterilisation techniques will also assist in reducing contamination issues. Replicate sampling across different DNA concentrations and mixed-individual contributions will help determine the marker recovery and sampling requirements for reliable identification at larger spatial scales. Nevertheless, our findings demonstrate the potential of targeted microhaplotype assays to extend eDNA applications from species detection to individual identification in tiger sharks, with potential applications to other low-template or degraded DNA sources, including museum and forensic samples (Barba et al., 2023; Solari et al., 2024). Ultimately, this approach could support large-scale monitoring of any large animals including marine megafauna, though success is likely dependent on species-specific ecology and DNA shedding dynamics (Barba et al., 2023).

## Acknowledgements

The authors acknowledge the Baiyungu, Yinigudira and West Thalanyji Traditional Owners of the Exmouth Gulf and neighbouring Nyinggulu (Ningaloo) coast and the Yamatji and Yued peoples of the Mid-West land and sea Country on which this research was conducted. Work on sea Country off Geraldton, Jurien Bay Marine Park and Exmouth Gulf was supported by Yamatji Southern Regional Cooperation Rangers, DBCA Baiyungu Rangers and DBCA Marine Rangers, respectively. We acknowledge collaborative support of Minderoo Foundation, the Minderoo Exmouth Research Laboratory and Minderoo OceanOmics Centre at UWA in this work and its mission to protect and restore natural ecosystems for future generations. Tiger shark icons incorporated throughout the manuscript are © marinewise.com.au. This project was partially funded with grants from the Ecological Society of Australia Holsworth Wildlife Research Endowment and the Jock Clough Marine Foundation via the Robson and Robertson Award.

## Data and code availability

The 55-marker multiplex primer information is available under Supplementary Materials 2. The sampling metadata, final microhaplotype and metabarcoding data are available under Supplementary Materials 3. Raw microhaplotype, metabarcoding and whole genome sequencing data will be available at a later date on NCBI/ENA.

## Ethics and permit approval statement

Samples from were collected under The University of Western Australia’s Animal Ethics Committee (approval numbers 2021/ET000634 and 2023/ET000935), fishing exceptions (numbers: 251157423, 251208424 and 251345625) granted by the Department of Primary Industries and Regional Development (DPIRD) and Fauna Taking and Lawful Authority Licence #FO25000591 Department of Biodiversity Conservation and Attractions (DBCA).

## Author contributions

J.R.P, L.T and S.C conceptualised and designed the study. S.C provided research funding. J.R.P, O.J, B.D and M.M undertook sample collection, while O.J, B.D, L.F and M.G.M provided the fieldwork funding and telemetry data. J.R.P and A.M conducted all lab work and data analysis, with S.C and L.T for data interpretation. J.R.P drafted the manuscript. All authors contributed to data interpretation and manuscript writing.

