## Supplementary materials for "Environmental DNA identifies individual free-ranging tiger sharks from a multi-use nuclear genotyping panel"

#### Section 1: Microhaplotype panel

**Table S1.** First-stage multiplex PCR and second-stage index PCR reagent volumes (µL). Tissue-derived gDNA was normalised to 2 ng/µL with 2 ng used as PCR input.

| **First-stage multiplex PCR** | | | **Second-stage index PCR** | |
|---|---|---|---|---|
| **Reagent** | **Sample type** | | **Reagent** | **Volume** |
| | gDNA | eDNA | | |
| 2X Qiagen Multiplex PCR Mastermix | 25 | 25 | Qiagen HotstarTaq Mastermix | 12.5 |
| Multiplex primer stock | 10 | 10 | Index primer | 1.5 |
| water | 14 | 10 | water | 10 |
| Tissue DNA | 1 | 5 | PCR product | 1 |
| Total reaction volume | 50 | 50 | Total reaction volume | 25 |

**Table S2.** First-stage multiplex PCR (two-step touchdown) cycling conditions.

| **#** | **Step** | **Temperature** | **Time** | **Number of Cycles** |
|---|---|---|---|---|
| 1 | Pre-Incubation | 95C | 15 minutes | 1 |
| 2 | Denaturation | 95C | 30 seconds | 15 |
| 3 | Annealing | 60C (-1C decrement) | 90 seconds | |
| 4 | Extension | 72C | 60 seconds | |
| 6 | Denaturation | 95C | 30 seconds | 30 |
| | Annealing | 56C | 90 seconds | |
| | Extension | 72C | 60 seconds | |
| 7 | Final extension | 72C | 10 minutes | 1 |
| 8 | Cooling | 4C | 10 minutes | 1 |

**Table S3.** Second-stage index PCR cycling conditions.

| **#** | **Step** | **Temperature** | **Time** | **Number of Cycles** |
|---|---|---|---|---|
| 1 | Pre-Incubation | 95C | 15 minutes | 1 |
| 2 | Denaturation | 95C | 30 seconds | 8 |
| 3 | Annealing | 62 | 30 seconds | |
| 4 | Extension | 72C | 30 seconds | |
| 7 | Final extension | 72C | 10 minutes | 1 |
| 8 | Cooling | 4C | 10 minutes | 1 |

#### Section 2: Bioinformatics

**Table S4.** Microhaplotype (MH) marker recovery per sample type after mapping and quality filtering. These reads were then used to call microhaplotypes.

| **Sample type** | **Mapped reads per MH/sample** | | **N MH/sample** | |
|---|---|---|---|---|
| | Mean | SD | Mean | SD |
| **Tissue** | 1786.03 | 2445.38 | 41.67 | 0.913 |
| **Paired eDNA** | 1773.14 | 862.19 | 41.76 | 0.59 |
| **Non-target eDNA** | 105.99 | 1892.58 | 13.88 | 17.21 |
| **Laboratory controls** | 0 | 0 | 0 | 0 |
| **Field controls** | 340.08 | 943.63 | 25.45 | 18.86 |

### Section 3: Analysis results

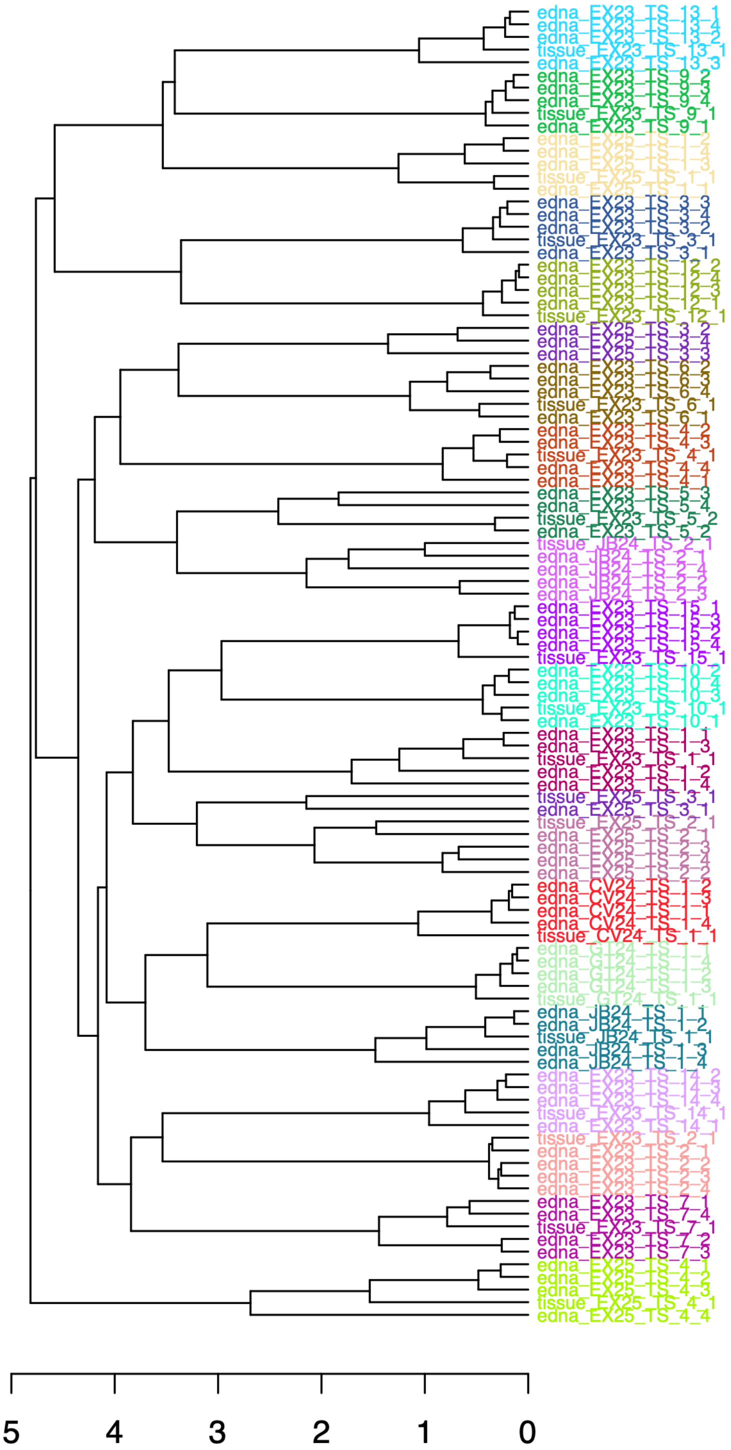

**Figure S1.** Dendrogram constructed using Euclidean genetic distances displaying the clustering between eDNA replicates and tissue samples from the same individual shark.

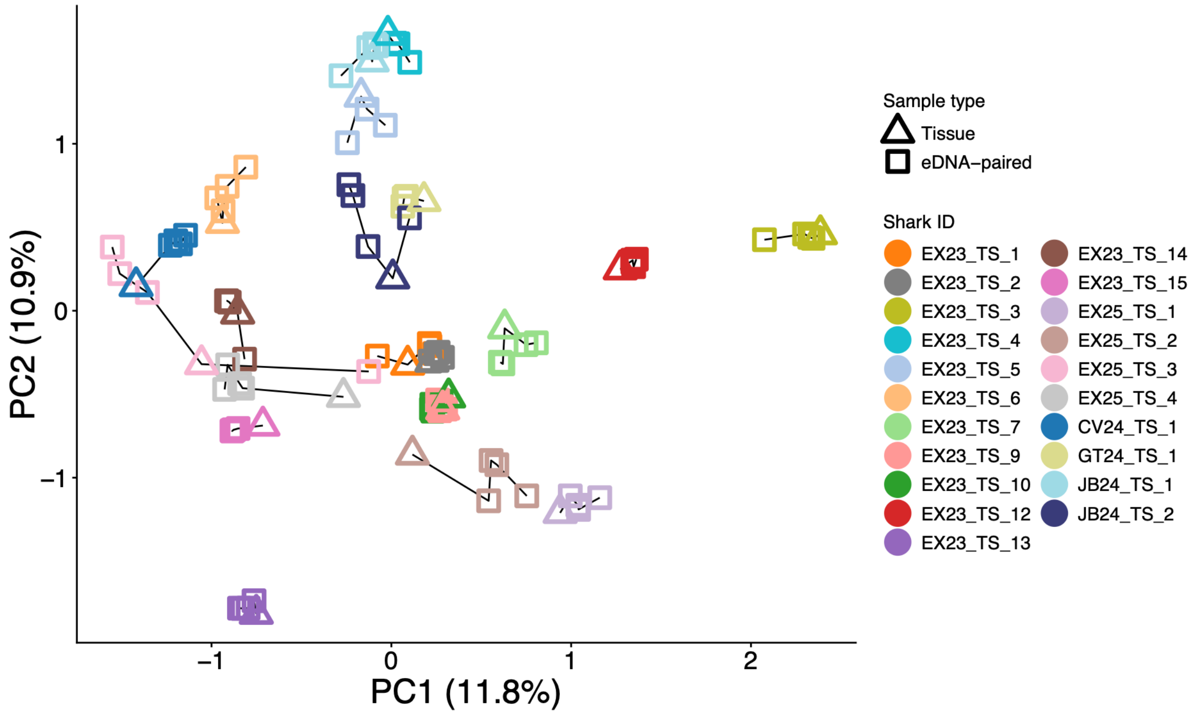

**Figure S2.** Principal Component Analysis (PCA) plot colour-coded by individual sharks, with solid black lines connecting sample pairs that correspond to the same individual.

**Table S5.** Number of SNPs filtered for population genetic comparison across whole genome sequencing data (WGS), microhaplotypes derived from tissue (MH-tissue) and eDNA samples (MH-eDNA), from the same 21 individual tiger sharks. Distance based thinning was conducted on all three datasets, but at different appropriate thresholds (WGS = 10kb; MH = 250bp). Linkage Disequilibrium (LD) pruning (r2 = 0.2) was also conducted as an alternate method (from the same filtered VCF file for each).

| Dataset | Called SNPs | Biallelic/ 10X | Filtered Mac=3 | Thinned | LD pruned |
|---|---|---|---|---|---|
| **Lc-WGS** | 12,782,851 | 9,251,452 | 5,725,383 | 282,010 | 631,696 |
| **MH-tissue** | 67 | 59 | 50 | 37 | 39 |
| **MH-eDNA** | 68 | 58 | 48 | 36 | 39 |

**Table S6.** Unbiased expected heterozygosity (uHe) and Inbreeding coefficient (FIS) comparison across whole genome sequencing data (WGS), microhaplotypes derived from tissue (MH-tissue) and eDNA samples (MH-eDNA), from the same 21 individual tiger sharks. Distance based thinning was conducted on all three datasets, but at different appropriate thresholds (WGS = 10kb; MH = 250bp). Linkage Disequilibrium (LD) pruning (r2 = 0.2) was also conducted as an alternate method (from the same filtered VCF file for each).

| **Dataset** | **uHe** | | | **FIS** | | |
|---|---|---|---|---|---|---|
| | Baseline | Thinned | Pruned | Baseline | Thinned | Pruned |
| **Lc-WGS** | 0.313 | 0.308 | 0.256 | 0.006 | 0.010 | -0.003 |
| **MH-tissue** | 0.325 | 0.337 | 0.341 | 0.084 | 0.099 | 0.036 |
| **MH-DNA** | 0.323 | 0.338 | 0.304 | 0.144 | 0.210 | 0.165 |

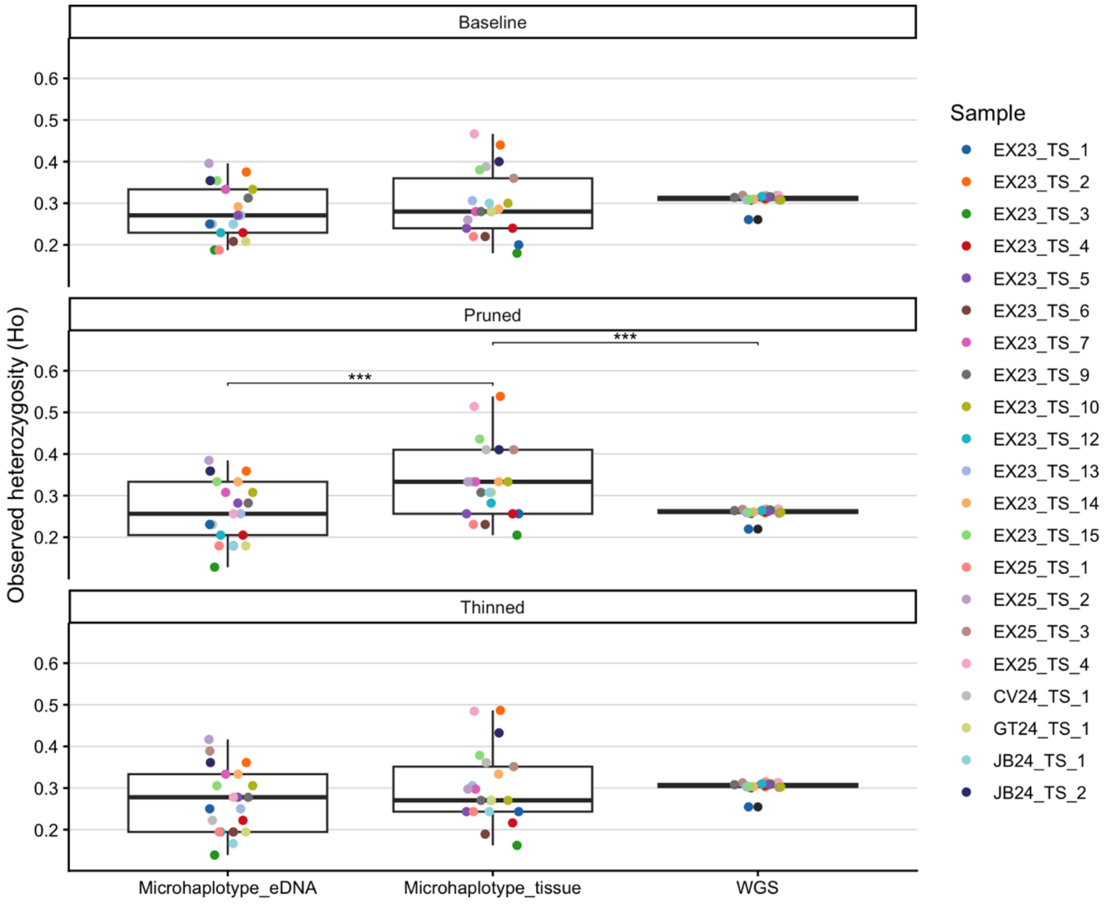

**Figure S3.** Observed heterozygosity (Ho) comparison across whole genome sequencing data (WGS), microhaplotypes derived from tissue (MH-tissue) and eDNA samples (MH-eDNA), from the same 21 individual tiger sharks. Distance based thinning was conducted on all three datasets, but at different appropriate thresholds (WGS = 50kb; MH = 250bp). Linkage Disequilibrium (LD) pruning (r2 = 0.2) was also conducted as an alternate method (from the same filtered VCF file for each).

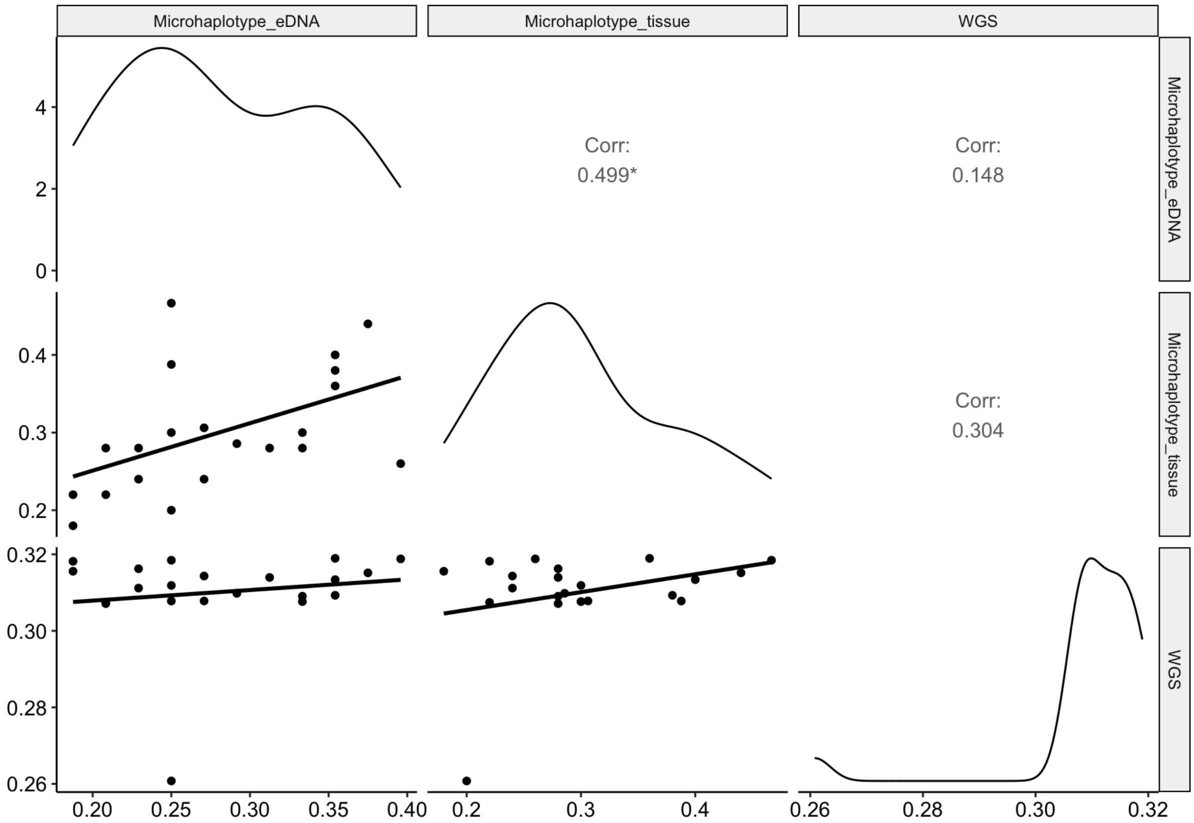

**Figure S4.** Pearson's correlation of Observed Heterozygosity (Ho) in individual samples between whole genome sequence (WGS) and microhaplotype data derived from both tissue and seawater eDNA. In this dataset, linkage disequilibrium (LD) was not accounted.

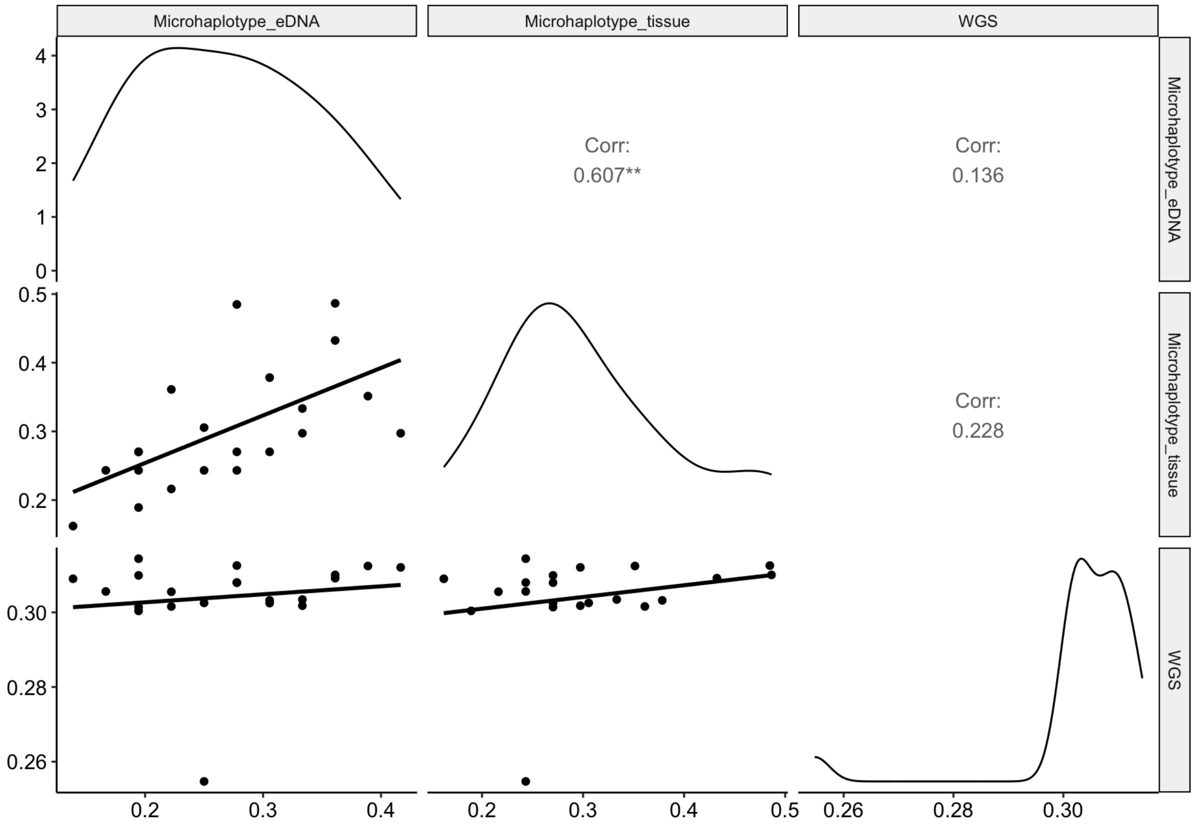

**Figure S5.** Pearson's correlation of Observed Heterozygosity (Ho) in individual samples (SNPs thinned) between whole genome sequence (WGS) and microhaplotype data derived from both tissue and seawater eDNA. In this dataset, linkage disequilibrium (LD) was accounted for using a distance-based thinning method.

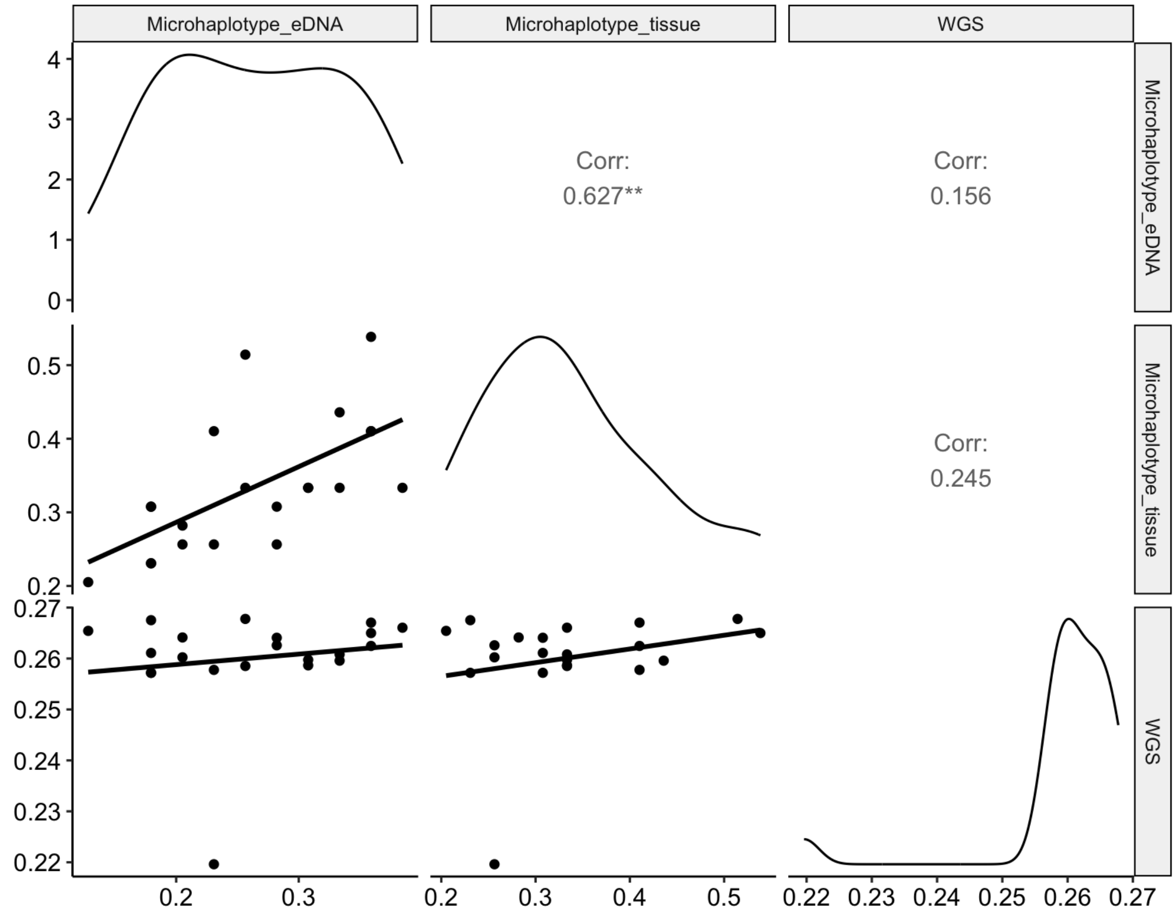

**Figure S6.** Pearson's correlation of Observed Heterozygosity (Ho) in individual samples (SNPs pruned) between whole genome sequence (WGS) and microhaplotype data derived from both tissue and seawater eDNA. In this dataset, linkage disequilibrium (LD) was accounted for using a pruning method (r2=0.2).

**Table S7.** Candidate identifications for non-targeted eDNA samples.

| Site | Marker recovery | Top candidate | | Second candidate | | Margin |
|---|---|---|---|---|---|---|
| | | ID | Similarity | ID | Similarity | |
| **CV24_CV** | 33 | tissue_CV24_TS_1 | 0.885 | tissue_JB24_TS_1 | 0.784 | 0.102 |
| **EX23_BS** | 31 | tissue_EX23_TS_12 | 0.897 | tissue_EX25_TS_2 | 0.798 | 0.0988 |
| **EX23_CS** | 6 | tissue_EX23_TS_6 | 0.738 | tissue_GT24_TS_1 | 0.731 | 0.007 |
| **EX23_LS** | 17 | tissue_EX23_TS_9 | 0.779 | tissue_EX23_TS_13 | 0.765 | 0.015 |
| **EX24_CS** | 2 | tissue_JB24_TS_1 | 0.887 | tissue_GT24_TS_1 | 0.822 | 0.065 |
| **GT24_GTS** | 37 | tissue_GT24_TS_1 | 0.932 | tissue_EX23_TS_12 | 0.768 | 0.164 |

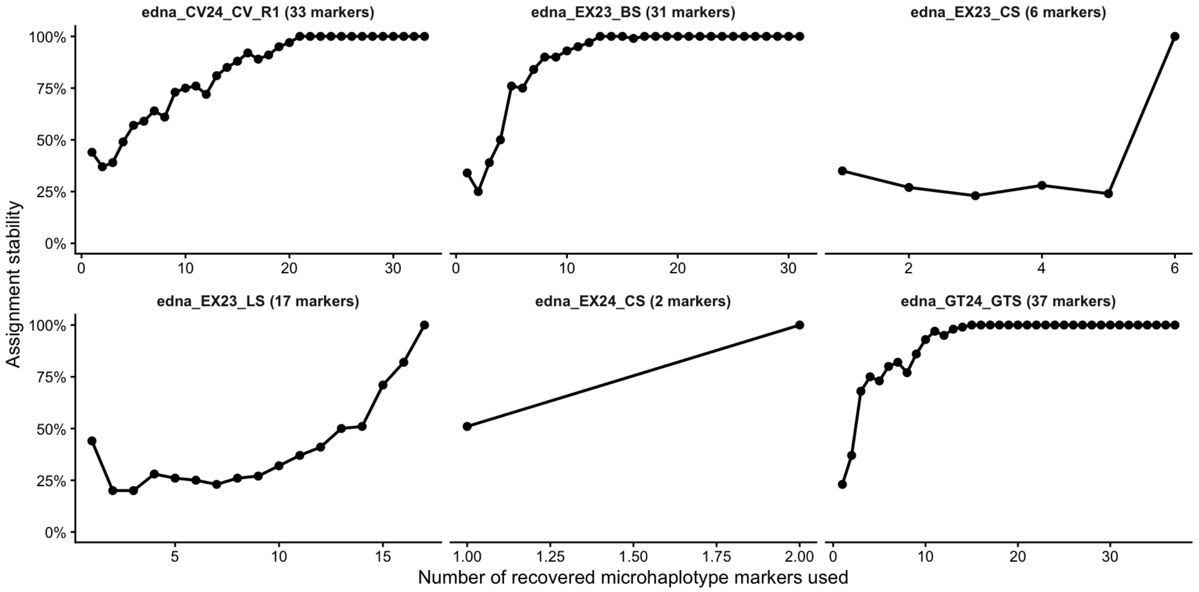

**Figure S7.** Assignment stability in non-target eDNA samples. Markers were randomly subsetted up to the maximum number recovered for each sample (ranged 2-37) for 100 bootstraps at each marker number. Assignment stability = proportion of same individual assigned for each marker number.

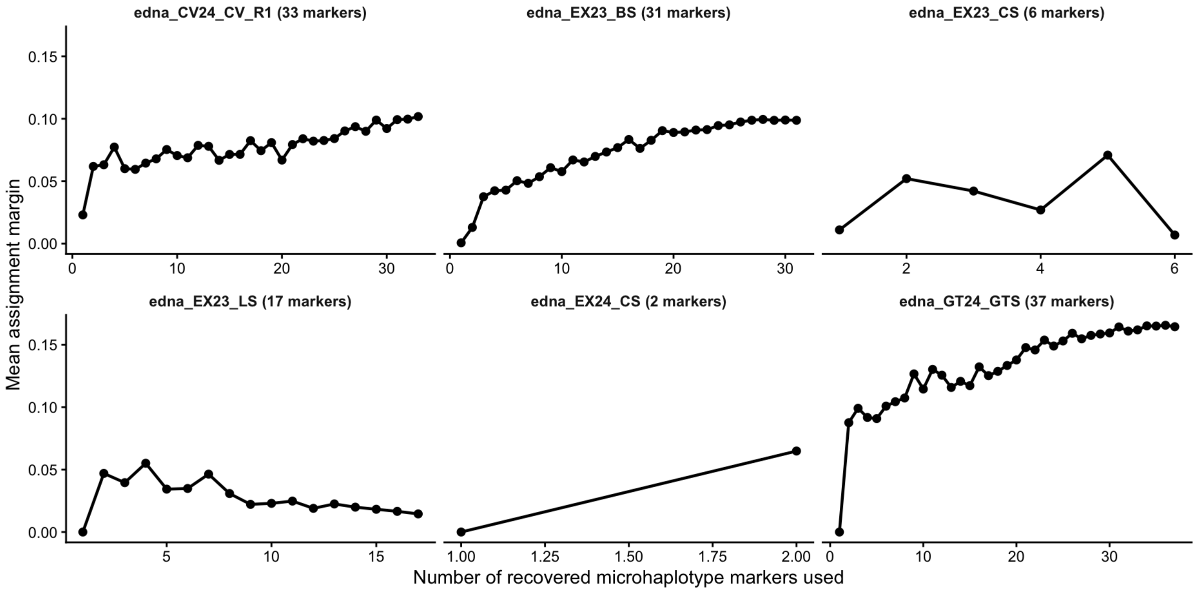

**Figure S8.** Assignment margin in non-target eDNA samples. Markers were randomly subsetted up to the maximum number recovered for each sample (ranged 2-37) for 100 bootstraps at each marker number. Assignment margin = difference in similarly score between the top and second-best candidate.
